# Leveraging mRNAs sequences to express SARS-CoV-2 antigens in vivo

**DOI:** 10.1101/2020.04.01.019877

**Authors:** Chunxi Zeng, Xucheng Hou, Jingyue Yan, Chengxiang Zhang, Wenqing Li, Weiyu Zhao, Shi Du, Yizhou Dong

## Abstract

SARS-CoV-2 has rapidly become a pandemic worldwide; therefore, an effective vaccine is urgently needed. Recently, messenger RNAs (mRNAs) have emerged as a promising platform for vaccination. Here, we systematically investigated the untranslated regions (UTRs) of mRNAs in order to enhance protein production. Through a comprehensive analysis of endogenous gene expression and de novo design of UTRs, we identified the optimal combination of 5’ and 3’ UTR, termed as NASAR, which was five to ten-fold more efficient than the tested endogenous UTRs. More importantly, NASAR mRNAs delivered by lipid-derived nanoparticles showed dramatic expression of potential SARS-CoV-2 antigens both in vitro and in vivo. These NASAR mRNAs merit further development as alternative SARS-CoV-2 vaccines.

## Introduction

SARS-CoV-2, a novel coronavirus, is causing a global pandemic, leading to over 900,000 confirmed cases and 45,000 death as of April 1, 2020^1^. SARS-CoV-2 shares 79% sequence identity with SARS-CoV and infects host cells through the receptor of angiotensin-converting enzyme 2^2^. To counteract this severe viral infection, a wide variety of therapeutics are in preclinical and clinical studies^3^. Currently, there is no FDA-approved treatment for SARS-CoV-2 infection.

Vaccine is the most effective strategy to prevent viral infections for a large population^4^. Different types of SARS-CoV-2 vaccines such as mRNA, DNA, and recombinant protein-based antigens are currently in clinical trials^5^. Among these agents, mRNA-based vaccine candidates first entered the clinical trial, because of the fast process for developing and manufacturing mRNA^6^. To express an antigen effectively, an mRNA requires several essential components, including 5’ cap, 5’ untranslated region (5’ UTR), antigen-encoding sequence, 3’ untranslated region (3’ UTR), and polyadenylated tail^7^. Among these components, the 5’ UTR and 3’ UTR are unique regulators for protein translation^8^. The design and selection of 5’ UTR and 3’ UTR are critically important to ensure the sufficient production of antigens and efficacious vaccination of the host^9^. Therefore, systematic explorations of UTRs may provide broad applicability for mRNA vaccines in response to emerging pathogens such as SARS-CoV-2. Previous studies utilized many UTRs from endogenous genes for protein expression^8, 10, 11^. For example, the UTRs from human cytochrome B-245 alpha polypeptide (CYBA) outperformed several other UTRs for protein expression^12^. The 5’ UTR from alpha globin mRNA enabled higher translation efficiency than that from beta globin mRNA in cells^13^. Alternatively, UTRs can be designed via a de novo method. A recent study reported a series of 5’UTRs consisting of 12-14 nucleotides (nt)^14^. Also, a deep learning method was applied to screen and analyze a large set of 5’UTRs^15^. Based on these significant advances, we hypothesize that the integration of endogenous UTRs with further de novo design may be a superior approach for UTRs engineering, facilitating the development of SARS-CoV-2 vaccines (**Fig. 1**). Our results prove the concept of this approach and significantly increase protein production compared to the UTRs reported in the literatures^12, 16-18^. Lastly, a series of potential SARS-CoV-2 antigens are visualized via fluorescent imaging in both cell and animal models, which may serve as vaccine candidates for the clinical trials.

**Figure 1.**
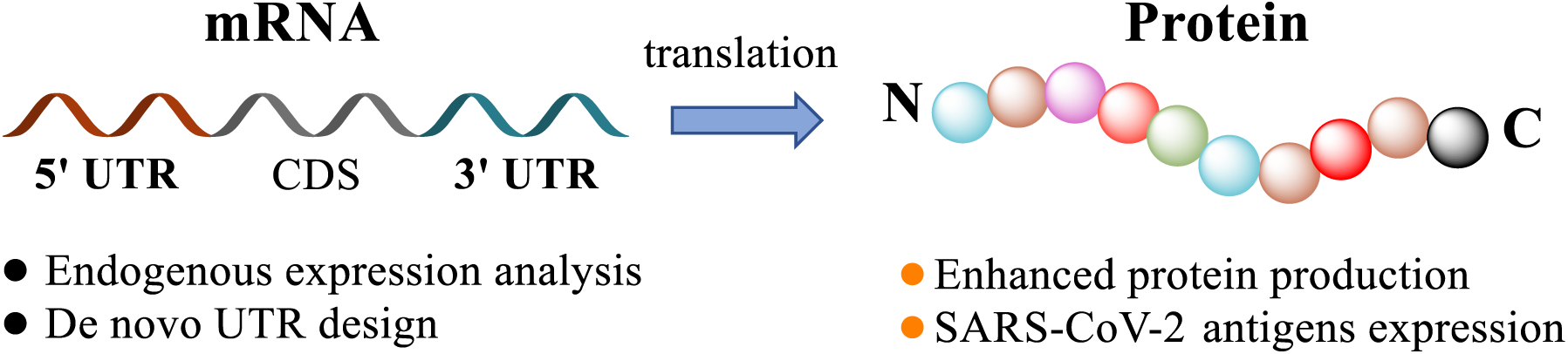
Schematic illustration of mRNA engineering. 5’ UTR and 3’ UTR of mRNA are comprehensively modulated based on the analysis of endogenous genes and further de novo design in order to enrich protein production and express SARS-CoV-2 antigens as potential vaccines.

## Results

### Investigation of endogenous 5’ UTR and 3’ UTR

#### Endogenous 5’ UTR from bioinformatics analysis

To screen endogenous UTRs in mammalian cells, we utilized a bioinformatics analysis of global gene expression as a starting point. Previously, the average half-life (h) and translation rate constant [protein copy number/(mRNA×h)] of 4248 mRNAs and corresponding proteins were quantified in mammalian cells^19^. According to their data, we calculated the translation capacity per mRNA molecule using the following equation: average half-life (h) × translation rate constant [protein copy number/(mRNA×h)] × protein length (amino acids) = translation capacity per mRNA molecule (total amino acids/mRNA) (**Supplementary Table 1**). These results indicated the number of amino acids produced by a single mRNA during its half-life. The mRNA from the murine Rps27a gene was found to possess the highest translation capacity per mRNA molecule. Rps27a is a housekeeping gene that encodes one of the components of the ribosome 40S subunit^20^. Two protein-coding transcripts exist for the murine Rps27a gene (named S27a-44 and S27a-45). These two transcripts have different 5’ UTRs and share the same 3’ UTR (**Supplementary Table 2** and **Supplementary Table 3**). Therefore, we generated two mRNAs utilizing S27a-45 and S27a-44 5’ UTRs, respectively. In these two 5’ UTRs, we found putative terminal oligo-pyrimidine (TOP) motifs consisting of 4-15 clustered pyrimidines (C and U), which were reported to negatively regulate mRNA translation (**Supplementary Table 2**)^21^. Subsequently, we removed the TOP motifs from S27a-44 and S27a-45 5’ UTRs and constructed two additional 5’ UTRs: S27a-44’ and S27a-45’. The mRNAs encoding Firefly luciferase were synthesized with these 5’ UTRs and were delivered to Hep3B and 293T cells using lipofectamine 3000. The control UTRs include CYBA UTRs (named CYBA)^12^, alpha globin UTRs (named AG)^13^, and modified alpha globin UTRs (AG+G with a complete Kozak sequence and 5AG+G without 3’ UTR)^18^. 5’ UTR from S27a-44 was better than that from S27a-45 (**Fig. 2a**). Removal of the putative TOP motif from S27a-45 moderately improved expression, while S27a-44’ without the TOP motif increased the relative luciferase activity over 70% compared to the S27a-44 in the two cell lines. S27a-44’ was comparable to CYBA and AG+G, the two best controls (**Fig. 2a**).

**Figure 2.**
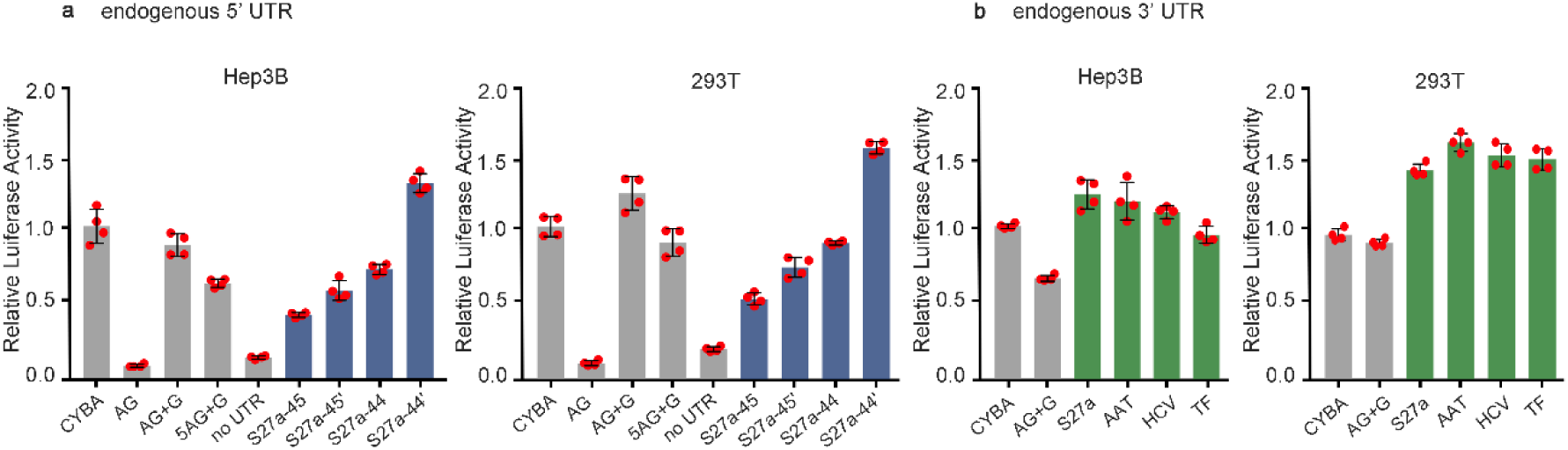
Evaluation of endogenous 5’ and 3’ UTRs. **a**, Relative luciferase activity in Hep 3B and 293 T cells. Four different 5’ UTR in blue are assessed with the same 3’ UTR. **b**, Relative luciferase activity in Hep 3B and 293T cells. Four different 3’ UTR in green are assessed with the same 5’ UTR. Control UTRs are in grey. Relative luciferase activity was normalized to that of CYBA. All data are presented as the mean ± s.d. (n=4).

#### Comparison study of endogenous 3’ UTR

We next compared 3’ UTR of the S27a with three alternative UTRs. Two 3’ UTRs were obtained from the mRNAs of transferrin (TF) and α-1-antitrypsin (AAT), two abundant human plasma proteins^22^. Another 3’ UTR from Hepatitis C viral RNA (HCV) was reported to retain and recycle the 40S subunit of ribosome after translation termination at stop codon, accelerating re-initiation of translation^23^. We then constructed these 3’ UTRs with the same 5’ UTR. As shown in **Fig. 2b**, these UTRs displayed similar effects on luciferase expression. Additionally, replacing S27a 3’UTR with another unstructured 3’ UTR of the same length significantly decreased protein expression (**Supplementary Table 3** and **Supplementary Fig. 1**). Since the length of S27a 3’ UTR (34nt) was considerably shorter than other 3’ UTRs (AAT 100nt, TF 386nt, or HCV 190nt), S27a 3’ UTR was used in the following studies.

### De novo design and optimization of 5’ UTR

#### Explorations of 5’ UTR length

Besides utilizing UTRs derived from endogenous genes, one alternative approach is to design mRNA UTRs via a de novo method, which is widely applied in protein engineering^24^. During the process of de novo design for UTRs, several general principles can be considered. For example, the Kozak sequence (GCCACC) located directly upstream of the translation start codon (AUG) can be included because it is a conserved sequence among many mammalian mRNAs for accurate start codon recognition^25^. Second, AUG within 5’ UTR is excluded to avoid alternative translation initiation and mutation of the amino acid sequence^10^. Third, secondary structures may be minimized to reduce the energy barriers for smooth scanning of the translation initiation complex along the 5’ UTR^13, 26^. Based on the knowledge in the literature, we designed 5’ UTR of different length: 10nt, 30, 50nt, 70nt, and 90nt with minimal secondary structures as modeled by RNAfold, a well-established computational tool for the prediction of RNA secondary structures (**Supplementary Fig. 2a** and **Supplementary Table 2**). We then prepared the mRNAs and evaluated their luciferase expression as mentioned above. The results showed that relative luciferase activity increased with 5’ UTR length and peaked at 70nt (**Fig. 3a**). Further increase of the length lowered luciferase expression. Therefore, 70nt was used as the optimal length for the subsequent design of 5’UTRs.

**Figure 3.**
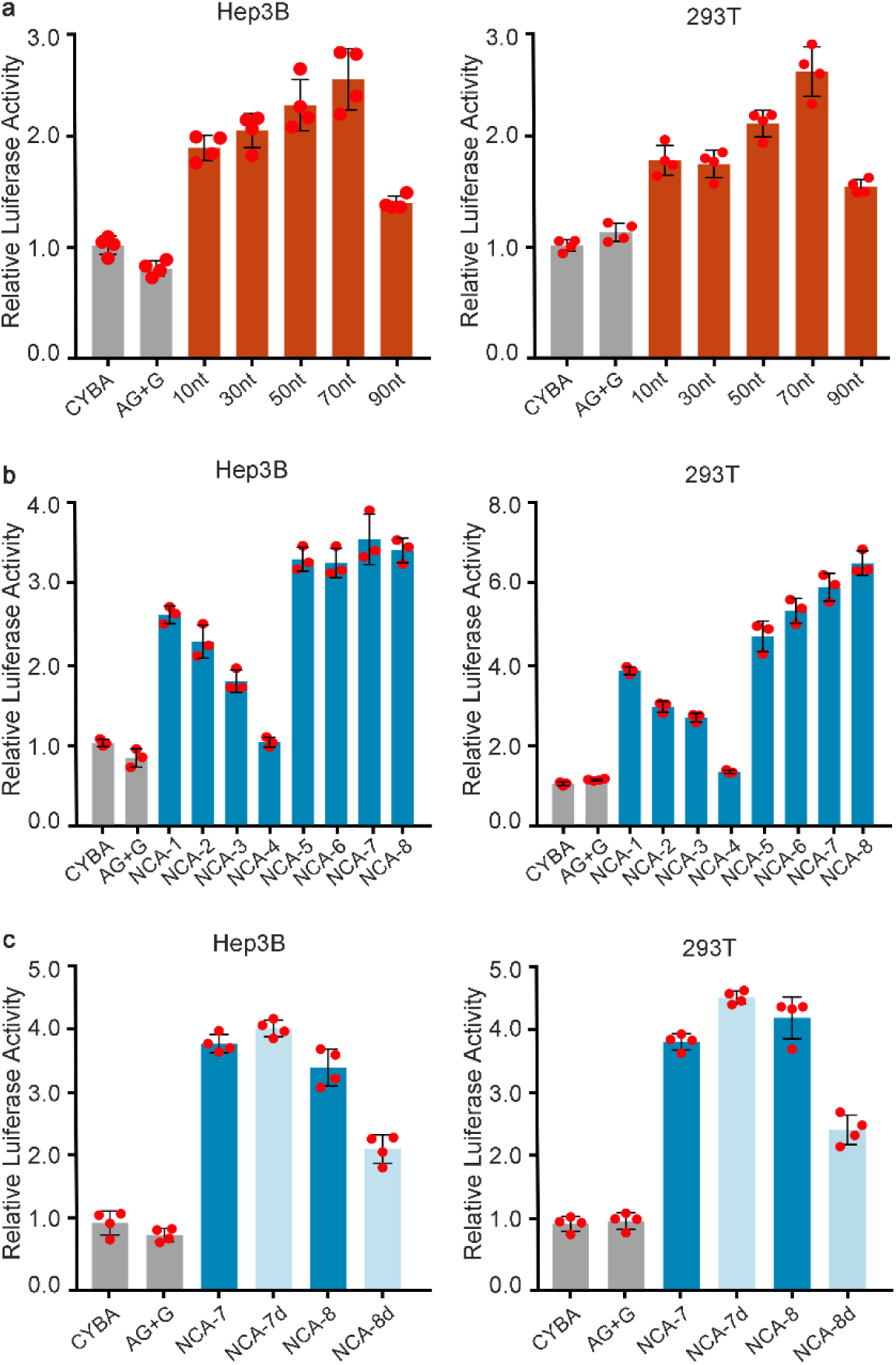
De novo design and engineering of 5’ UTR. **a**, Relative luciferase activity of mRNAs with 5’ UTRs consisting of 10nt, 30nt, 50nt, 70nt, or 90nt in Hep 3B and 293T cells. **b**, Relative luciferase activity of mRNA with different nucleotide compositions of 5’ UTRs in Hep 3B and 293T cells. **c**, Relative luciferase activity after removal of miRNA target sites in 5’ UTR. NCA-7d and NCA-8d: removal of miRNA binding sites from NCA-7 and NCA-8, respectively. Relative luciferase activity was normalized to that of CYBA. All data are presented as the mean ± s.d. (n=4).

#### Nucleotide composition analysis of 5’ UTR

In addition to length, nucleotide compositions of A, C, G, and U may be an important factor at the 5’ UTR. Generally, high GC content is commonly associated with increased structural complexity^27^. G can base pair with both C and U and induce the formation of more secondary structures^27^. The increase of U content may introduce additional chemically modified nucleotides into 5’ UTR if modified U, such as pseudouridine are used in mRNA synthesis. To understand the effects of nucleotide compositions, we performed nucleotide compositions analysis and designed a series of 5’ UTRs (NCA-1 to NCA-8). When adjusting nucleotide composition, the minimal numbers of G and C are 3 and 4, respectively, due to the transcription start site determined by the T7 promoter sequence (GG) and the use of Kozak sequence (GCCACC). These 5’ UTRs were also built to avoid secondary structures (**Supplementary Fig. 2b**), thereby minimizing the impact of structures on translation initiation. As shown in **Fig. 3b**, a decrease of relative luciferase activity was observed with less A and more G when the amount of C and U were constant (NCA-1 vs NCA-2; NCA-1 vs NCA-3). When G and U were minimal, the increase of C and decrease of A content showed a moderate decrease of luciferase expression (NCA-3 vs NCA-4). When holding G content to the minimal of 3, the inclusion of U boosted relative luciferase activity (NCA-5 vs NCA-6 vs NCA-7 vs NCA-8). Moreover, these 5’ UTR with ACU outperformed those consisting of mostly AC and AG, suggesting the importance of U on 5’ UTR. Consequently, two constructs with high U contents, NCA-7 and NCA-8, were selected for further studies.

#### Removal of microRNA target sites from 5’ UTR

MicroRNAs were previously considered to mainly target the 3’ UTR of mRNA for gene regulations^28^. However, recent data showed that miRNAs also interfered with ribosomal scanning by binding to target sites within 5’ UTR^29^. Hence, we examined possible miRNA target sites within NCA-7 and NCA-8 using the prediction tool available at miRDB. The prediction exhibited that both NCA-7 and NCA-8 contained miRNA target sites (**Supplementary Table 2**). Then, we adjusted the order of the nucleotides of NCA-7 and NCA-8 and created NCA-7d and NCA-8d, which maintain their nucleotide compositions with minimal secondary structure. Interestingly, NCA-7d showed a slightly increased expression than NCA-7, while NCA-8d was less active than NCA-8 (**Fig. 3c**). Based on these results, NCA-7d was selected as the lead 5’ UTR.

#### Optimization of 3’ UTR with integrated motifs

Next, we further studied the effects of functional motifs integrated with the lead S27a 3’ UTR. Previous studies reported that abundant cis-regulatory elements existed in mRNA 3’ UTR to modulate protein expression, such as functional motifs for RNA binding proteins (RBPs)^11^. Specific RBPs are able to bind to their cognate motifs within mRNA 3’ UTR and enhance protein expression through various mechanisms^30^. One of the RBPs, QKI-7, was shown to enable cytoplasmic polyadenylation of mRNA and upregulation of protein expression after binding to its target motif called QKI Response Element (QRE) in 3’ UTR^31^. Two QRE sequences were obtained from previous reports, namely QRE1^32^ and QRE2^33^. Another RBP, HuR, was believed to stabilize target transcripts by binding to its cognate motifs called AU-Rich Element (ARE) in eukaryotic cells^34^. One such ARE sequence was obtained from human eIF4E mRNA 3’ UTR^34^. Besides eukaryotic mRNA, the sindbis virus (SinV) evolved to utilize part of its viral 3’ UTR to recruit HuR and induce potent stabilization of the viral RNA genome^35^. The SinV RNA regions responsible for HuR binding, including repeated sequence element 3 and U-rich element (together named R3U) was obtained. Additionally, the ribosome binding fragments from two internal ribosome entry sites (IRESs) of encephalomyocarditis virus (EMCV)^36^ and foot-and-mouth disease virus (FMDV)^37^ were selected. These two fragments possess high binding affinities to ribosomal subunits and have the potential to retain and recycle the subunits for more efficient translation re-initiation. Based on these literature findings, we prepared mRNA transcripts with the integration of the following elements after S27a 3’ UTR: QRE1, QRE2, R3U, ARE, EMCV, and FMDV (**Supplementary Table 3**). R3U was identified as a preferred functional motif from the results in both Hep3B and 293T cells (**Fig. 4a**). Through the comprehensive UTR engineering, the optimal UTRs was identified as NCA-7d as the 5’ UTR and S27a plus a functional motif R3U as the 3’ UTR (named as NASAR). After several rounds of design and validation of UTRs, we then compared NASAR with S27a-45, the starting point of endogenous gene together with two additional control UTRs (MOD1 and MOD2) in the literature^16, 17^. NASAR was over 10-fold more potent than S27a-45, 4 to 7-fold better than CYBA or AG+G, and up to 2-fold superior to MOD1 and MOD2 in cell lines and primary cells (**Fig. 4b** and **Supplementary Fig. 3**). These results demonstrate that our concept, integration of endogenous UTRs with further de novo design, is an efficient strategy for UTRs design.

**Figure 4.**
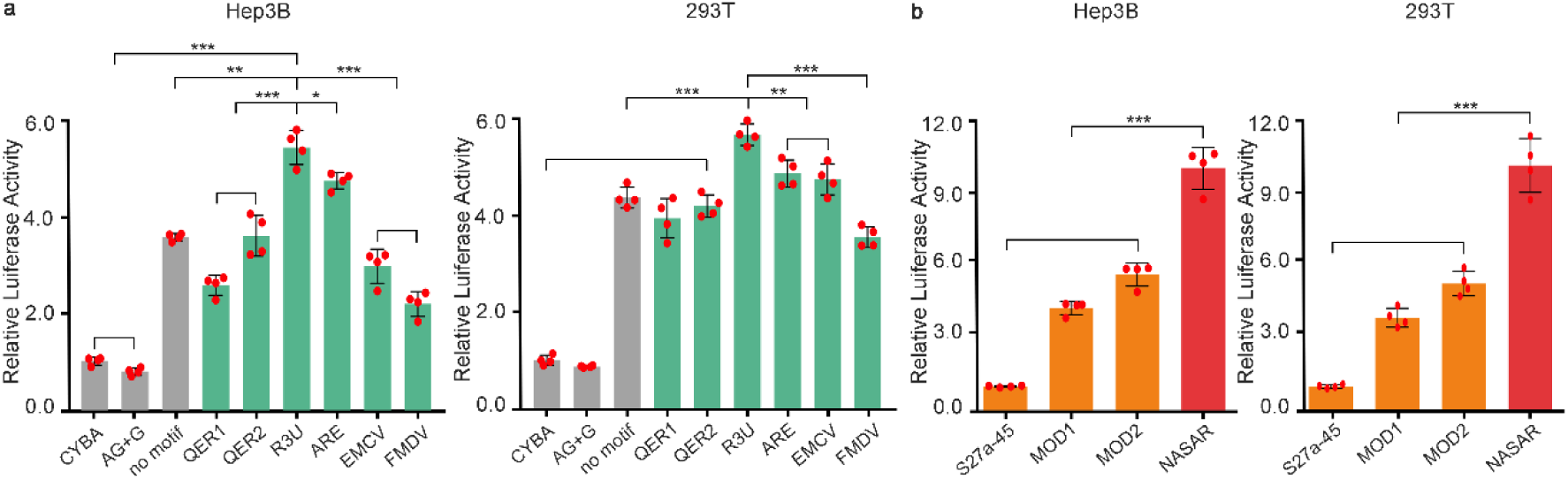
Modifications of 3’ UTR. **a**, Relative luciferase activity of mRNAs with addition of RNA motifs after S27a 3’ UTR. No motif: S27a 3’ UTR only. All engineered mRNAs in green utilized the same 5’ UTR. **b**, Relative luciferase activity of NASAR mRNAs in comparison to S27a-45, a start UTR, and additional control UTRs, MOD1 and MOD2. Relative luciferase activity was normalized to that of S27a-45. All data are presented as the mean ± s.d. (n=4). Statistical significance in a and b was analyzed by the two-tailed Student’s t-test. \**P* < 0.05; \*\**P* < 0.01; \*\*\**P* < 0.001; n.s., not significant.

#### NASAR mRNAs express SARS-CoV-2 antigens

Given the potent activity of NASAR mRNAs and urgent demand for SARS-CoV-2 vaccines, we aim to create NASAR mRNAs encoding SARS-CoV-2 antigens. According to previous studies on SARS-CoV, its structural spike (S) and nucleocapsid (N) proteins, regarded as the dominant antigens eliciting neutralizing antibodies in SARS-CoV patients, were explored as vaccine candidates^38, 39^. The receptor binding domain (RBD) of the S protein responsible for viral entry was also utilized for vaccine development^38^. The membrane (M) protein contained epitopes^40^. Additionally, the envelope (E) protein interacting with M was indispensable for the assembly of viral particles^41^. Therefore, five NASAR mRNAs were prepared to express RBD, S, N, M, and E proteins as the vaccine candidates. Since no antibodies have been developed against these five potential antigens in SARS-CoV-2, we installed a FLAG tag to RBD andS protein, and a VSV-G tag to N, M, and E proteins in order to detect their expression. To confirm the effects of UTRs on antigen expression using NASAR mRNA vaccine, three additional mRNAs were synthesized using the coding sequence of RBD with control UTRs, CYBA, MOD2, and NASAR. After the delivery of these three mRNAs into 293T cells using our previously developed lipid like-nanoparticles, TT3^42^, RBD expression was analyzed by an ELISA. NASAR enabled antigen expression 3-fold of CYBA and 1.6-fold of MOD2 (**Fig. 5a**), consistent with the luciferase expression results in **Fig. 4c**. Then, NASAR mRNAs encoding all five SARS-CoV-2 antigens were delivered to 293T cells by TT3 separately. We observed the obvious expression of these antigens through immunostaining (**Fig. 5b, Supplementary Fig. 4**).

**Figure 5.**
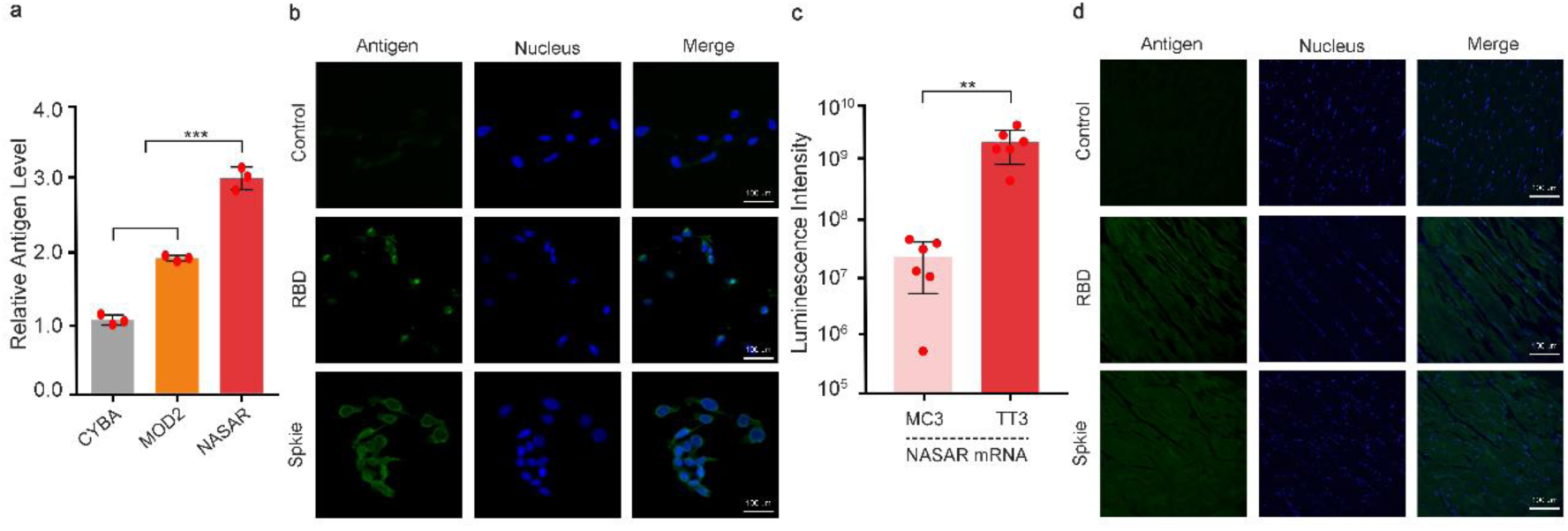
NASAR mRNAs encoding potential SARS-CoV-2 antigens. **a**, Relative RBD antigen level in 293T cells quantified by ELISA (n=3). **b**, Fluorescent microscopy imaging of S and RBD antigens in 293T cells. Scale bar = 100µm. **c**, Quantification of luciferase expression in vivo after i.m. injection to mice. MC3-NASAR luciferase and TT3-NASAR luciferase mRNAs (n=6). **d**, Confocal microscopy imaging of S and RBD antigens expressed in mouse muscle tissues after i.m. injection. Scale bar = 100µm. All data are presented as the mean ± s.d. Statistical significance in a and c was analyzed by the two-tailed Student’s t-test. \*\**P* < 0.01; \*\*\**P* < 0.001.

Based on the encouraging cell results, we performed intramuscular (i.m.) injection, a widely used administration route for vaccines^43^, of NASAR mRNAs. Similar to in vitro results, NASAR was 4.2-fold more potent than CYBA in the luciferase expression when both were delivered by TT3 nanoparticles (**Supplementary Fig. 5**). Moreover, TT3 nanoparticles induced over 70-fold more luciferase activity than MC3 nanoparticles, an FDA-approved delivery vehicle^44, 45^, when formulated with the same NASAR mRNAs (**Fig. 5c**). Lastly, NASAR mRNAs encoding either RBD or S protein were formulated with TT3 and injected intramuscularly into hind legs of mice. Confocal microscopy imaging of the mouse tissues with immunostaining confirmed the expression of both antigens (**Fig. 5d**).

## Discussion

This study describes a rational engineering approach to enhance protein production using mRNA. Through bioinformatics analysis and modification of endogenous 5’ UTR and 3’ UTR, followed by de novo design and further optimization, the most effective NASAR UTR was identified, enabling strong expression of multiple potential SARS-CoV-2 antigens both in vitro and in vivo.

Based on this study, several criteria should be considered in the design of mRNA UTRs: (1). Regarding the 5’ UTR, the length is critical and the optimal sequences tested in this work do not exceed 70nt. And it should be free of certain regulatory elements, e.g. TOP motifs, secondary structure, upstream open reading frame, and microRNA binding site. (2). In terms of 3’ UTR, secondary structures may benefit protein production. A plausible explanation is that such structure in 3’ UTR might prevent ribosome from skipping the stop codon and reaching mRNA 3’ end that triggers Non-Stop mRNA Decay (NSD)^46^. Long 3’ UTR length does not necessarily provide additional translation benefits. Long 3’ UTR typically contains a mixture of positive and negative cis-regulatory elements^11^. Although it is challenging to obtain a long endogenous 3’ UTR with only positive elements, specific positive cis-regulatory motifs in 3’ UTRs may be used to improve mRNA translation. R3U, a HuR binding motif added to 3’ UTR, enhanced protein expression, consistent with the observation in their natural context^34, 35^. This indicates that certain positive cis-regulatory elements may be used as an important UTR component.

In order to cope with the current SARS-CoV-2 pandemic, NASAR mRNAs can be applied to encode various SARS-CoV-2 antigens. Importantly, TT3 formulated NASAR Fluc mRNA was much more efficient for than an FDA-approved delivery system MC3. This enables us to combine the strength of NASAR mRNAs and TT3 nanoparticles to maximize the expression of SARS-CoV-2 antigens. Overall, this work provides a proof-of-concept study to develop mRNA vaccine candidates against SARS-CoV-2, which may be conducive to clinical translations.

## Methods

### Chemicals and reagents

Opti-MEM Reduced Serum Medium was purchased from Thermo Fisher Scientific. Cell culture plates and luminescent assay plates were obtained from Corning. MC3 was purchased from MedKoo. DOPE and DMG-PEG2000 were purchased from Avanti Polar Lipids. Cholesterol was obtained from Sigma-Aldrich. Q5 High-Fidelity PCR Kit was purchased from NEB.

### De novo design of 5’ UTR

5’ UTRs were designed de novo using AntaRNA (http://rna.informatik.uni-freiburg.de) ^47^ with predefined nucleotide composition as the sequence constrain and minimal secondary structure as the structural constrain. GU base pairs were permitted. The secondary structure of the output sequences was further confirmed by RNAfold (http://rna.tbi.univie.ac.at/cgi-bin/RNAWebSuite/RNAfold.cgi). Sequences with undesired secondary structures were discarded. The microRNA binding sites within 5’ UTR were predicted using the custom prediction tool in the microRNA database (miRDB) (http://mirdb.org/)^48^. Due to the minimum size limit of 100nt for mRNA target submission, the full 5’ UTR and first 100 nucleotides of the firefly luciferase coding region were used as the input sequences.

### Plasmid construction

The sequences necessary for in vitro transcription of mRNA, including T7 promoter, 5’ UTR, coding sequence, and 3’ UTR, were cloned into pUC19 vector using repliQa HiFi Assembly Mix (QuantaBio) and transformed into 5-alpha Competent E. coli (NEB) by chemical transformation. The transformed E. coli was allowed to grow on LB broth (Miller) plate with agar and 100ug/mL carbenicillin (Teknova). Individual colonies were inoculated and outgrown in LB broth liquid medium (Miller) containing 100ug/mL carbenicillin overnight with vigorous shaking at 250 rpm. Plasmids were extracted using QIAprep Spin Miniprep Kit (Qiagen). Concentration was measured on a NanoDrop 2000 Spectrophotometer (Thermo). The region of interest in the plasmid from the T7 promoter to 3’ UTR was confirmed by Sanger Sequencing.

### mRNA synthesis

All mRNA transcripts were synthesized by in vitro transcription using a protocol modified from our previous publication^49^. Briefly, DNA templates were synthesized by PCR amplification of the corresponding plasmids using a forward primer and a reverse primer containing 120T at 5’ end. The DNA templates were purified by QIAquick PCR Purification Kit (Qiagen) and examined by agarose gel electrophoresis. All mRNAs were synthesized by in vitro transcription with 100% substitution of UTP by pseudouridine-5’-triphosphate (TriLink) using AmpliScribe T7-Flash Transcription Kit (Lucigen) following the manufacturer’s instruction and purified by RNA Clean & Concentrator (Zymo). The capping of mRNA was conducted using Vaccinia Capping System (NEB) and Cap 2’-O-Methyltransferase (NEB), followed by another purification by RNA Clean & Concentrator (Zymo). After measurement of concentration by a NanoDrop 2000 Spectrophotometer (Thermo), all mRNAs were diluted to the desired concentration in 1× TE, aliquoted, and stored at -80°C for future use.

### Firefly luciferase assay in vitro

Hep3B and 293T cells were cultured in Eagle’s Minimum Essential Medium (Corning) with 10% Fetal Bovine Serum (FBS) and Dulbecco’s Modified Eagle Medium (Corning) with 10% FBS, respectively. Hep3B and 293T cells were seeded at a density of 2×10^4^ cells/well on a white 96-well flat-bottom plate (Costar) followed by overnight incubation. The mRNAs were thawed on ice and denatured at 65°C for 3min followed by incubation at room temperature for 45min. When Lipofectamine 3000 (Thermo) was used, the transfection was performed according to the manufacturer’s instruction. When TT3 nanoparticles was used, mRNA was formulated using lipid:DOPE:cholesterol:DMG-PEG2000 at a molar ratio of 20:30:40:0.75 as reported before.^42^ Then, cells in each well were treated with 50 ng per well of Firefly luciferase mRNA. The luminescence activity was determined on a Cytation 5 Cell Imaging Multi-Mode Reader (Biotek) using a Bright-Glo Luciferase Assay Kit (Promega).

### Quantification of RBD levels with ELISA

1 × 10^5^ 293T cells were seeded in each well in a 24-well plate and grew overnight. The cells were treated with 100 ng per well mRNAs encoding the RBD antigen using TT3 nanoparticles as mentioned above. After overnight incubation, cells were lysed in each well. The cell lysis was retrieved and centrifuged at 12000×rpm for 10min. EZview Red ANTI-FLAG M2 Affinity Gel was used to concentrate the FLAG-tagged RBD in the pellet following the manufacturer’s protocol. RBD was eluted into 0.1M glycine HCl at pH 3.5 and diluted 10 times before coating a 96-well immunoplate (Thermo/Nunc). After blocking, primary rabbit anti-FLAG antibody (ab1162, abcam) at 1:1000 dilution was added, followed by incubation with HRP-linked anti-rabbit IgG (Cell Signaling, 7074). OD492 reading was obtained on a Cytation 5 Cell Imaging Multi-Mode Reader (Biotek).

### Immunostaining and microscopy imaging

293T cells were grown overnight on cover slides placed in a 6-well plate. TT3 nanoparticles were formulated with mRNAs encoding SARS-CoV-2 antigens, S, RBD, M, N, and E proteins, as described above. Six hours after treatment, cells were washed with phosphate buffered saline (PBS) and fixed with 10% formalin, followed by permeabilization with 0.2% Tween-20. Staining was conducted using primary rabbit anti-FLAG antibody (abcam, ab1162) at 1:200 dilution and FITC-linked secondary goat anti-rabbit polyclonal antibody (abcam, ab6717) at 1:1000 dilution. Nucleus was stained with Hoechst 33342 (Thermo). After sealing the slides, images were taken on a Nikon Eclipse Ti Inverted Fluorescence Phase Contrast Microscope with NIS-Elements BR imaging software (version 4.20).

### Luciferase expression assay in vivo

All mouse studies were approved by the Institutional Animal Care and Use Committee at The Ohio State University and complied with local, state, and federal regulations. mRNA encoding Firefly luciferase was formulated at the concentration of 0.03 mg/mL using TT3 nanoparticlesand dialyzed in 1×PBS. Three mice per group were injected intramuscularly 15 µg mRNA per leg (n=6 legs). Six hours after injection, mice were imaged using a Xenogen IVIS Spectrum In Vivo imaging system (Caliper)

### SARS-CoV-2 antigens expression in vivo

The mRNAs encoding SARS-CoV-2 antigens, RBD and S, were formulated by TT3 nanoparticles to 0.03 mg/mL and 0.06 mg/mL, respectively. The formulated nanoparticles were dialyzed in 1×PBS. The formulation was injected intramuscularly 15µg RBD-encoding mRNA or 30µg S-encoding mRNA. Mice were euthanized six hours after injection. The muscle tissue at the injection sites was harvested and incubated in 4% paraformaldehyde overnight, followed by 30% sucrose incubation overnight. The tissue was sectioned using Cryotome E Cryostat (Thermo-Shandon) and fixed using acetone. The subsequent staining procedure was the same as that for staining 293T cells. Images were taken on a Nikon A1R Live Cell Confocal microscope with NIS-Elements BR imaging software (version 4.20).

### Data analysis

All data analysis was conducted in Prism 7 (GraphPad). All t-tests were two-tailed and *P < 0.05, **P < 0.01, ***P < 0.001, ****P < 0.0001 was considered statistically significant. The p values and specific statistical methods were shown in the figure legends.

## Acknowledgements

We acknowledge the OSU Campus Microscopy & Imaging Facility for providing the instruments and services. Y.D. acknowledges the support from the National Institutes of Health (NIH) through the Maximizing Investigators’ Research Award R35GM119679 of the National Institute of General Medical Sciences as well as the start-up fund from the College of Pharmacy at Ohio State University.

## Author contributions

C.Zeng conceived, designed and performed the experiments, analyzed data, and wrote the paper. X.H. performed ELISA and animal experiments. J.Y. and W.L. contributed to tissue imaging. S.D., C. Zhang, and W.Z. contributed to animal experiments. J.Y contributed to plasmid construction. Y.D. conceived and supervised the project and wrote the paper. The final manuscript was approved by all authors.

## Competing interests

The authors declare no competing interests.

Correspondence and requests for materials should be addressed to Y.D.

