## Supplementary information for "Leveraging mRNAs sequences to express SARS-CoV-2 antigens in vivo"

**Supplementary Table 1**. Translation capacity per mRNA molecule of 4248 genes. The genes were sorted based on their translation capacity per mRNA. The 30 genes with the highest translation capacity were list below. The full dataset is available in the Additional Information file. The information and data of Gene Names, protein length [amino acids], average mRNA half-life [h], and average translation rate constant [molecules/(mRNA*h)] are adapted from the literature [1]. Translation capacity per mRNA (total amino acids/mRNA) is calculated using the following equation: protein length (amino acids) × average half-life (h) × translation rate constant (protein copy number/(mRNA×h)).

| Gene Names | Protein length [amino acids] | Average mRNA half-life [h] | Average translation rate constant [molecules/(mRNA*h)] | Translation capacity per mRNA (total amino acids/mRNA) |
|---|---|---|---|---|
| Rps27a | 156 | 10.26 | 224528.75 | 359371736 |
| Hist1h1c | 212 | 3.85 | 147337.15 | 120256582 |
| H2afv | 129 | 19.97 | 29040.14 | 74811176 |
| Rab11b | 218 | 15.3 | 19290.92 | 64342935 |
| Dnaja1 | 397 | 11.31 | 12921.59 | 58018844 |
| Naca | 2187 | 14.15 | 1869.6 | 57856735 |
| Hnrnpd | 355 | 22.13 | 5169.69 | 40613860 |
| Rpl15 | 204 | 9.04 | 20242.96 | 37331257 |
| Hmgb2 | 210 | 10.11 | 17120.18 | 36347854 |
| Rpl31 | 125 | 9.86 | 25219.48 | 31083009 |
| Ahcy | 432 | 15.84 | 4414.95 | 30210973 |
| Tep1 | 2629 | 14.78 | 764.43 | 29703166 |
| Dync1h1 | 4644 | 23.58 | 270.23 | 29591677 |
| Serbp1 | 407 | 31.92 | 2155.29 | 28000321 |
| Ncl;Nuc | 707 | 27.85 | 1394.12 | 27450153 |
| Plec1 | 4691 | 10.2 | 548.39 | 26239474 |
| Psma3 | 255 | 12.33 | 8076.35 | 25393256 |
| Rpl18a | 176 | 21.5 | 6465.83 | 24466701 |
| Tln1 | 2541 | 24.99 | 383.78 | 24369873 |
| Cse1l | 971 | 11.13 | 2247.8 | 24292492 |
| Rps15a | 130 | 5.3 | 27988.42 | 19284021 |
| Dpysl2 | 572 | 6.05 | 5327.25 | 18435481 |
| Myo9a | 2631 | 6.7 | 1029.41 | 18146131 |
| Elavl1 | 326 | 9.44 | 5683.23 | 17489799 |
| Ywhaz | 245 | 11.11 | 6288.32 | 17116493 |
| Cad | 2225 | 26.89 | 273.81 | 16382121 |
| Flnb | 2602 | 20.05 | 307.7 | 16052740 |
| Pdia3 | 505 | 25.68 | 1232.02 | 15977328 |
| Lmna | 665 | 18.79 | 1277.63 | 15964434 |

**Supplementary Table 2**. Sequences of 5’ UTRs used in this study.

| Name | Sequence | Annotation | Source |
|---|---|---|---|
| S27a-45 | GGGAGGAAAGCCUCUCUUAAUCGC<br>AUCGGCUGUAUAAGAAAGCCUUUU<br>GAGGCAUUUUUUUUAGUUGAGCAC<br>AUCAUUUCGAGGCCAUUCUGAGGU<br>AAACCGAGAAAAGAGCGUAAAGAA<br>ACCGAGCGAACGAGCAAAUCUGGC<br>ACUGCGUUAGACAGCCGCGAUUCCG<br>CUGCAGCGCGCAGGCACGUGUGUG<br>GCCGCCUAAGGGGCGGGUCCUUCGG<br>CCAGGAGACCCCGUCGGCCACGCUC<br>GGAUCUUCCUUUCCGAUCCGCCAUC<br>GUGGGUGGAGCCGCCGCCACG | underline: putative TOP motif | Ensembl transcript ID ENSMUST000 00102845.10 |
| S27a-45’ | GGGAGGAAAGAAUCGCAUCGGCUG<br>UAUAAGAAAGCCUUUUGAGGCAUU<br>UUUUUUAGUUGAGCACAUCAUUUC<br>GAGGCCAUUCUGAGGUAAACCGAG<br>AAAAGAGCGUAAAGAAACCGAGCG<br>AACGAGCAAAUCUGGCACUGCGUU<br>AGACAGCCGCGAUUCCGCUGCAGCG<br>CGCAGGCACGUGUGUGGCCGCCUAA<br>GGGGCGGGUCCUUCGGCCAGGAGA<br>CCCCGUCGGCCACGCUCGGAUCUUC<br>CUUUCCGAUCCGCCAUCGUGGGUGG<br>AGCCGCCGCCACG | | modifed from S27a-45 |
| S27a-44 | GGGUUUCCGAUCCGCCAUCGUGGG<br>UGAGUGUAUGCUCUGUGGCCGCGC<br>UCUGGCUAGUGGCGCUACGCGUCGC<br>UCUCACGGGUGUCGUCGGAUCUAA<br>UCCGUCUCUUUUCGAAUGCAGGUG<br>GAGCCGCCGCCACG | underline: putative TOP motif | Ensembl transcript ID ENSMUST000 00102844.3 |
| S27a-44’ | GGGGAUCCGCCAUCGUGGGUGAGU<br>GUAUGCUCUGUGGCCGCGCUCUGGC<br>UAGUGGCGCUACGCGUCGCUCUCAC<br>GGGUGUCGUCGGAUCUAAUCCGUC<br>UCUUUUCGAAUGCAGGUGGAGCCG<br>CCGCCACG | | modifed from S27a-44 |
| 10nt | GGGAGCCACC | | De novo design |
| 30nt | GGGAAAGAAACAGGACAGAAAACA<br>GCCACC | | De novo design |
| 50nt | GGGAACGACAAGAAACACAUACAA<br>AAGAAACAGGACAGAAAACAGCCA<br>CC | | De novo design |

| | | | |
|---|---|---|---|
| 70nt | GGGAAGAGAUAAACAUAAACAUAA<br>ACGACAAGAAACACAUACAAAAGA<br>AACAGGACAGAAAACAGCCACC | | De novo design |
| 90nt | GGGGAGAAGAGGGAACAGGACACA<br>AGAGAUAAACAUAAACAUAAACGA<br>CAAGAAACACAUACAAAAGAAACA<br>GGACAGAAAACAGCCACC | | De novo design |
| NCA-1 | GGACACAAACAAACAAACAAACAC<br>ACAAACACACAAACAAACACACACA<br>CACACAAACAAACAAGCCACC | | De novo design |
| NCA-2 | GGACACACACACACACACACACACA<br>CACACACACAAACACACACACACAC<br>ACACAAACACACAAGCCACC | | De novo design |
| NCA-3 | GGAAAGAGAGAAAGAGAGAAAGAA<br>AGAAAGAGAGAGAGAAAGAGAGAA<br>AGAAAGAAAGAGAGAAGCCACC | | De novo design |
| NCA-4 | GGAGAGAGAGAGAGAGAGAAAGAG<br>AGAGAGAGAGAAAGAGAGAGAGAG<br>AGAGAGAGAGAGAGAAGCCACC | | De novo design |
| NCA-5 | GGAAACACAUAAACAUAAACAUAA<br>CACACAACAAACACAUACAACACAA<br>ACACAACACAAAACAGCCACC | | De novo design |
| NCA-6 | GGAAACACAAUAACAUAAUCAUAC<br>UACACAACUAACACAUACAUCACAU<br>ACACAUCACAUAACAGCCACC | | De novo design |
| NCA-7 | GGAAACACAAUAACAUAAUCAUAC<br>UACACAACUAACACAUACAUCACAU<br>ACACAUCACAUAACAGCCACC | Underline: microRNA target sites | De novo design |
| NCA-8 | GGCUACACACUCUCACUCUCAUCAC<br>UCACUACUCACUCUCUCAUCACUCU<br>CACAUCACAUCACUGCCACC | Underline: microRNA target sites | De novo design |
| NCA-7d | GGCAAAAAUCAAAAUCAAUCAUCA<br>UCACAACAUCAACAAUCAAUCAUCA<br>ACACAUCAUCAAGACACCACC | | Derived from NCA-7 |
| NCA-8d | GGCAUCACACUCUCACUCUCAUCUC<br>AACACUCCUCCUCAUUCCAAUCUCU<br>CACACAUCCCAUUAGCCACC | | Derived from NCA-8 |

**Supplementary Table 3**. Sequences of 3’ UTRs used in this study. The miRNA target sites were predicted by miRDB[2], a well-established online tool[3].

| Name | Sequence | Annotation | Source |
|---|---|---|---|
| S27a | UUGUGUAUGCGUUAAUAAAAAG<br>AAGGAACUCGUA | | Ensembl transcript IDs: ENSMUST000<br>00102845.10 and ENSMUST000<br>00102844.3 |
| AAT | CUGCCUCUCGCUCCUCAACCCCU<br>CCCCUCCAUCCCUGGCCCCCUCCC<br>UGGAUGACAUUAAAGAAGGGUU<br>GAGCUGGUCCCUGCCUGCAUGUG<br>ACUGUAAA | | Ensembl transcript ID: ENST0000039<br>3087.8<br>Genbank ID NM_000295 |
| TF | AAUCUCAGAGGUAGGGCUGCCAC<br>CAAGGUGAAGAUGGGAACGCAG<br>AUGAUCCAUGAGUUUGCCCUGGU<br>UUCACUGGCCCAAGUGGUUUGUG<br>CUAACCACGUCUGUCUUCACAGC<br>UCUGUGUUGCCAUGUGUGCUGAA<br>CAAAAAAUAAAAAUUAUUAUUG<br>AUUUUAUAUUUCAAAAACUCCAU<br>UCUUUCCUAAAUAUUUUCAACAA<br>AGGAUUUCUUUAUGCAUUCUGCC<br>UAAAUACCUAUGCAACUGAGCCC<br>UUCCUUCUCAGCUCAAGAUUCGU<br>CUGGUCUUUCCCUACAGCUUUGU<br>GUGUGCCAUGGCCACAUCUCCUG<br>GGUACAGUUCAAGGAGACAUCUU<br>UUCUAAAAGGGUCUGCGUGAUCA<br>UUAAAAUAUAAUCAAAUGUA | | ENST000004<br>02696.7 |
| HCV | AGGUUGGGGUAAACACUCCGGCC<br>UCUUAAGCCAUUUCCUGUUUUUU<br>UUCUUUCCUUUCCUUCUUUUUUU<br>CCUUUCUUUUUCCCUUCUUUAAU<br>GGUGGCUCCAUCUUAGCCCUAGU<br>CACGGCUAGCUGUGAAAGGUCCG<br>UGAGCCGCAUGACUGCAGAGAGU<br>GCUGAUACUGGCCUCUCUGCAGA<br>UCAUGU | | [4] |
| unstructured | CAAUUUUCCGCUCGAGCUCUAUA<br>UGAUACAACAU | | De novo design |
| S27a + QRE1 | UUGUGUAUGCGUUAAUAAAAAG<br>AAGGAACUCGUA | underline: QRE1 | [5] |

| | | | |
|---|---|---|---|
| | GCCGUAACCACGUCUACUAACGCCG | | |
| S27a + QRE2 | UUGUGUAUGCGUUAAUAAAAAGAAGGAACUCGUA AACUCCAGGACUGUAUUUGUGACUAAUUGUAUAACAGGUU | underline: QRE2 | 6 |
| S27a + R3U | UUGUGUAUGCGUUAAUAAAAAGAAGGAACUCGUA AAAACUCAAUGUAUUUCUGAGGAAGCGUGGUGCAUAAUGCCACGCAGCGUCUGCAUAACUUUUAUUAUUUCUUUUAUUAAUCAACAAA | underline: R3U | 7 |
| S27a + ARE | UUGUGUAUGCGUUAAUAAAAAGAAGGAACUCGUA UAUGUCUGUUUUUGUAUCUUUAUGCUGUAUUUUAACACUUUGUAUUACUUAGGUUAUU | underline: ARE | 8 |
| S27a + EMCV | UUGUGUAUGCGUUAAUAAAAAGAAGGAACUCGUA AUAAGAUACACCUGCAAAGGCGGCACAACCCCAGUGCCACGUUGUGAGUUGGAUAGUUGUGGAAAGAGUCAAAUGGCUCUCCUCAAGCGUAUUCAACAAGGGGCUGAAGGAUGCCCAGAAGGUACCCCAUUGUAUGGGAUCUGAUCUGGGGCCUCGGUGCACAUGCUUUACAUGUGUUUAGUCGAGGUUAAAAAACGUCUAGGCCCCCCGAACCACGGGGACGUGGUUUUCCUUUGAAAAACACGAUGAUAAU | underline: EMCV IRES | 9 |
| S27a + FMDV | UUGUGUAUGCGUUAAUAAAAAGAAGGAACUCGUA CAGCAGGUUUCCCCAACUGACACAAAACGUGCAACUUGAAACUCCGCCUGGUCUUUCCAGGUCUAGAGGGGUAACACUUUGUACUUGGUGACAGGCUAAGGAUGCCCUUCAGGUACCCCGAGGUAACACGCGACACUCGGGAUCUGAGAAGGGGACUGGGGCUUCUAUAAAAGCGCUCGGUUUAAAAAGCUUCUAUGCCUGAAUAGGUGACCGGAGGUCGGCACCUUUCCUUUACAAUUAAUGACCCU | underline: FMDV IRES | 10 |

**Supplementary Table 4**. Sequences of the control UTRs used in this study.

| Name | 5' UTR | 3' UTR | Ref. |
|---|---|---|---|
| AG | GGGACUCUUCUGG UCCCCACAGACUCA GAGAGAACCCACC | GCUGGAGCCUCGGUGGCCAUGC UUCUUGCCCCUUGGGCCUCCCC CCAGCCCCUCCUCCCCUUCCUGC ACCCGUACCCCCGUGGUCUUUG AAUAAAGUCUGAGUGGGCGGCA | 11 |
| 5AG+G | GGGACUCUUCUGG UCCCCACAGACUCA GAGAGAACGCCACC | | 12 |
| AG+G | GGGACUCUUCUGG UCCCCACAGACUCA GAGAGAACGCCACC | GCUGGAGCCUCGGUGGCCAUGC UUCUUGCCCCUUGGGCCUCCCC CCAGCCCCUCCUCCCCUUCCUGC ACCCGUACCCCCGUGGUCUUUG AAUAAAGUCUGAGUGGGCGGCA | Modified from 12 |
| CYBA | GGGCGCGCCUAGCA GUGUCCCAGCCGGG UUCGUGUCGCC | CCUCGCCCCGGACCUGCCCUCCC GCCAGGUGCACCCACCUGCAAU AAAUGCAGCGAAGCCGGGA | 13 |
| MOD1 | GGGAAAUAAGAGA GAAAAGAAGAGUA AGAAGAAAUAUAA GAGCCACC | GCUGCCUUCUGCGGGGCUUGCC UUCUGGCCAUGCCCUUCUUCUC UCCCUUGCACCUGUACCUCUUG GUCUUUGAAUAAAGCCUGAGUA GGAAG | 14 |
| MOD2 | GGGAAAUAAGAGA GAAAAGAAGAGUA AGAAGAAAUAUAA GAGCCACC | UGAUAAUAGGCUGGAGCCUCGG UGGCCAUGCUUCUUGCCCCUUG GGCCUCCCCCCAGCCCCUCCUCC CCUUCCUGCACCCGUACCCCCGU GGUCUUUGAAUAAAGUCUGA | 15 |

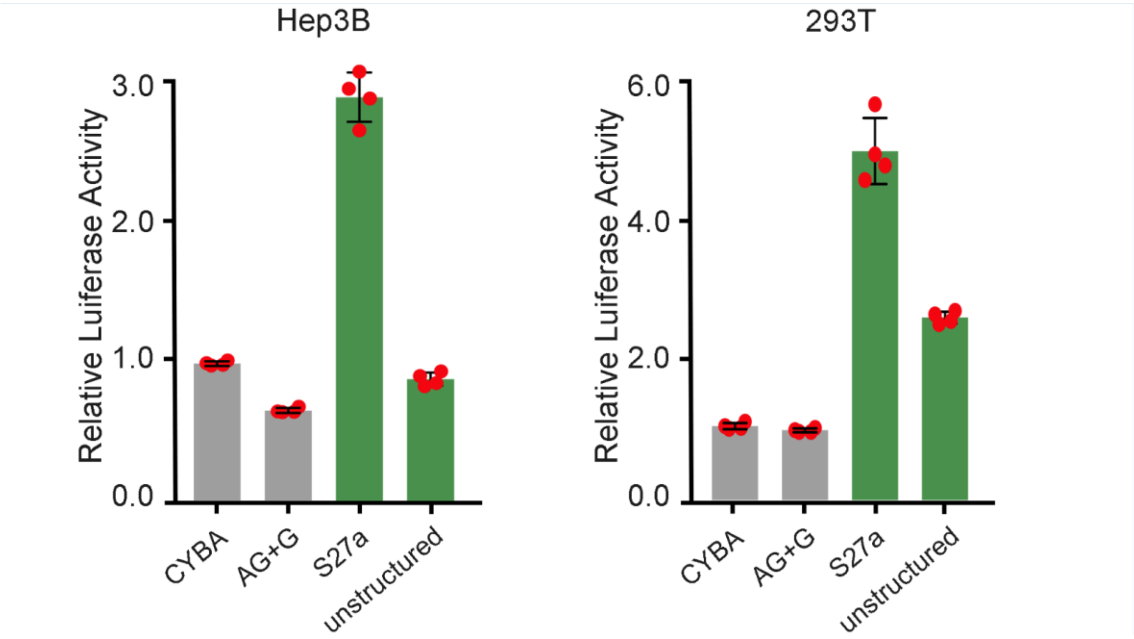


**Supplementary Figure 1.** Modification of 3’ UTR. Relative luciferase activity in Hep 3B and 293T cells. Two different 3’ UTR in green are assessed with the same 5’ UTR. Relative luciferase activity was normalized to that of CYBA. All data are presented as the mean ± s.d. (n=4).

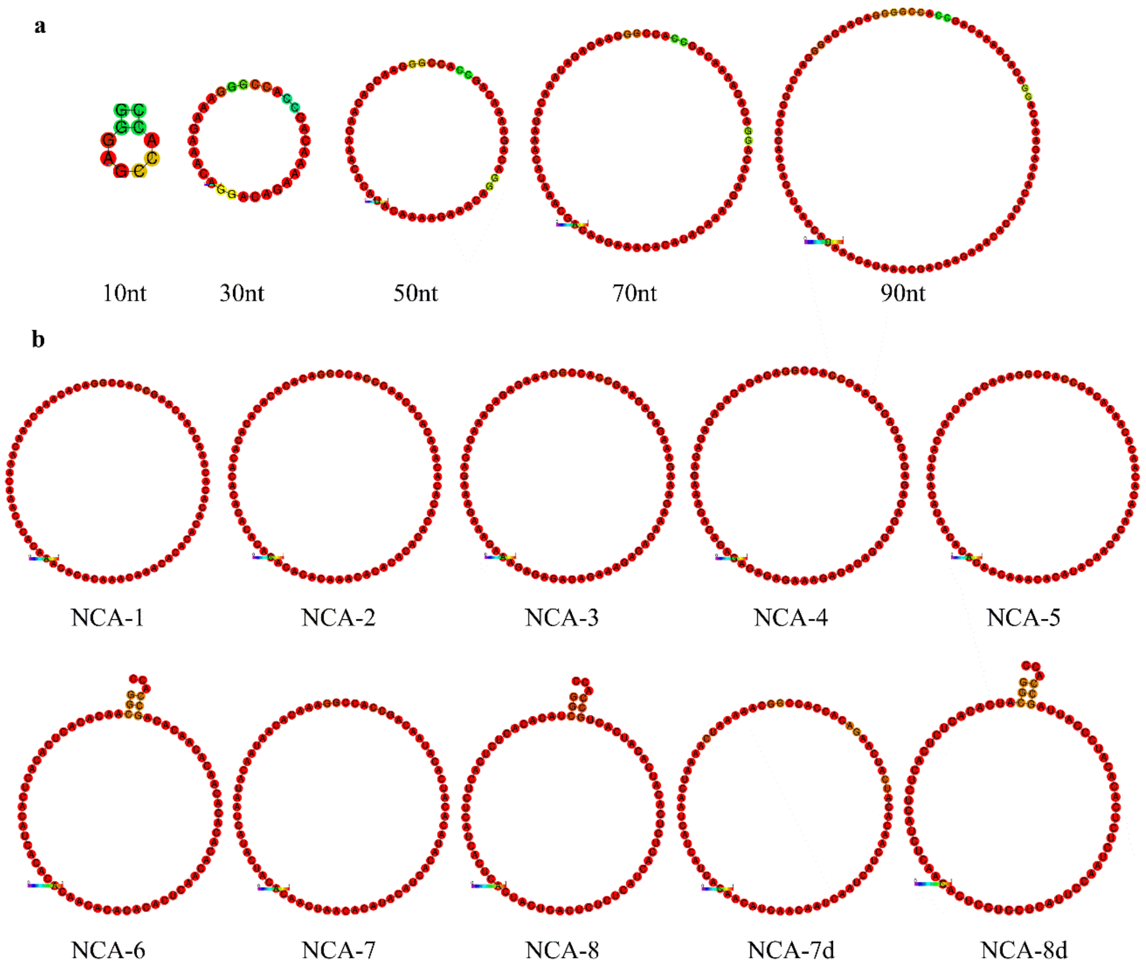


**Supplementary Figure 2.** Secondary structures prediction of de novo designed 5' UTRs. **a**, 5'UTRs of various lengths. **b**, 5' UTRs designed from nucleotide composition analysis. Minimum free energy (MFE) secondary structures were predicted by RNAfold, a well-established online tool[16]. Colors indicate base-pair probabilities with red and green showing high and low probability, respectively.

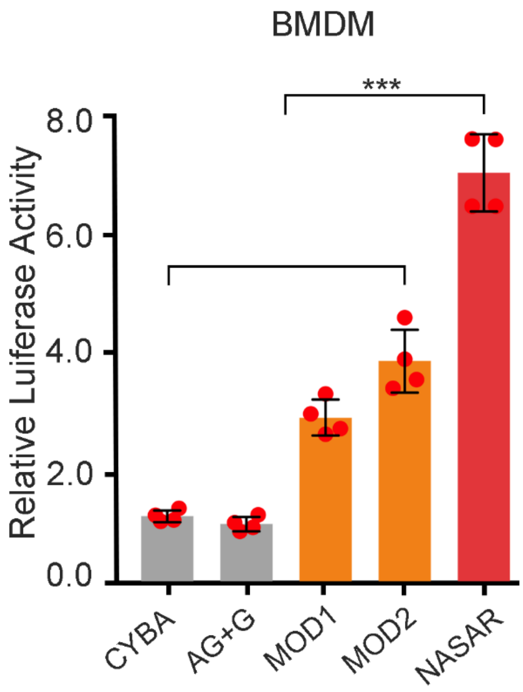


**Supplementary Figure 3.** Evaluation of NASAR mRNA in primary cells. Relative luciferase activity in bone marrow derived macrophages (BMDM). Relative luciferase activity was normalized to that of CYBA. All data are presented as the mean ± s.d. (n=4). Statistical significance in a and c was analyzed by the two-tailed Student's t-test. ***$P$ < 0.001.

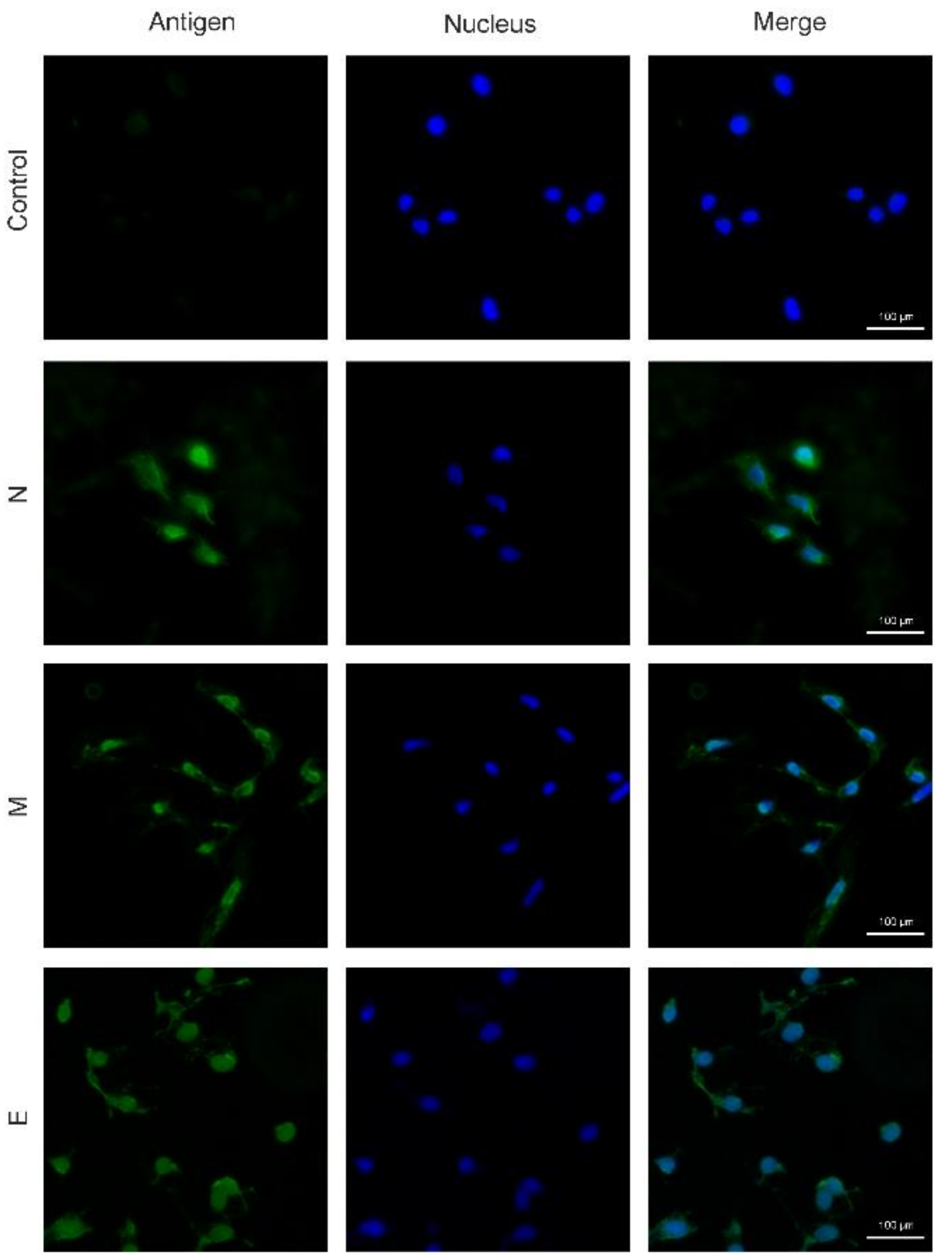


**Supplementary Figure 4.** NASAR mRNAs expressing SARS-CoV-2 antigens. Fluorescent microscopy imaging of Nucleocapsid (N), Membrane (M), and Envelope (E) proteins in 293T cells. Scale bar = 100µm.

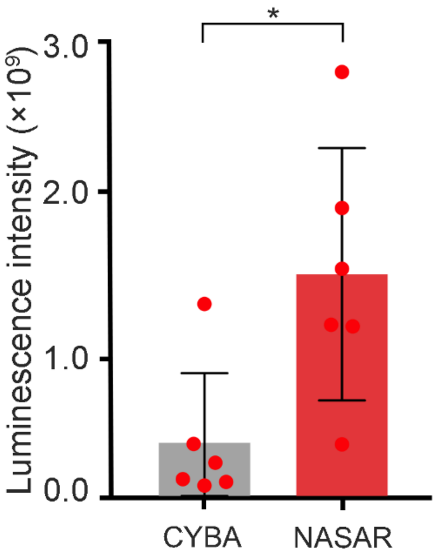


**Supplementary Figure 5.** Quantification of luciferase expression in vivo after i.m. injection to three mice per group (n=6 legs). Both mRNAs were formulated with TT3 nanoparticles. All data are presented as the mean ± s.d. Statistical significance in a and c was analyzed by the two-tailed Student's t-test. $^*P < 0.05$.
